# Postural perturbation induced by electrical stimulation; A new approach to examine the mechanism of falling

**DOI:** 10.1101/2021.03.25.437084

**Authors:** Behdad Tahayori, Bahman Tahayori, Alireza Mehdizadeh, David M. Koceja

## Abstract

**Background:** Falling is a major cause of disability and death among elderly people. Therefore, a clear understanding of fall mechanism is necessary for providing preventative and treatment methods. Several fall simulation protocols have been introduced to study lost of balance in a laboratory setting.

**New Method:** We have explained and examined a new method to induce a sudden perturbation on standing posture to provide an insight into the mechanism of falling. The method comprises eliciting an H-reflex protocol while subjects are standing which produces a contraction in soleus and gastrocnemius muscles. We have also defined analytical techniques to provide biomarkers of balance control during perturbation. The method is easy to implement and interpret. The H-reflex or M-wave can be elicited unilaterally or bilaterally causing a forward or sideway perturbation. The vector analysis and the Equilibrium Point calculations defined here can quantify the amplitude, direction, and evolution of the perturbation.

**Results:** We tested this method on a group of healthy individuals and observed clear patterns of loss of balance due to stimulation. Direction and magnitude of deviation was manifested through the reconstructed vectors, with bilateral stimulation causing the largest perturbation.

**Comparison and conclusion:** The resultant plantarflexion torque is reminiscent of tripping over an obstacle and triggers corrective reactions to restore balance. Therefore, it is more similar to an internal perturbation. Mechanical perturbations to the torso cause a displacement in center of mass (COM) and trigger a cascade of mechanisms. Our method, does not trigger the perturbation by the displacement of COM initially and therefore, triggers fewer mechanisms for regaining balance.

## Introduction

Loss of balance and over ground fall is a major cause of disability and death among elderly people in the United States (Galet et al. 2018, Kerber et al. 1998). Elderly people’s falls are notoriously dangerous; up to 60% of falls in elderly population result in severe physical injuries. Recent years statistics show that more than 50% of fall patients are being placed in long-term care and nursing facilities (Voermans et al. 2007). However, replicating an actual fall in a laboratory environment is challenging because it is and involuntary and - most of the time - unpredictable event. Nonetheless, understanding the neural and mechanical events preceding a fall and the events immediately after the initiation of loss of balance (LOB) are critical to understand the underlying causes/mechanisms of fall. Therefore, although it is impossible to record a naturally occurring fall in a lab, it is possible to induce a mechanical LOB to observe the critical events before and immediately after the LOB. Unfortunately, the mechanism of perturbation determines the type of responses and events leading to LOB and fall (Bortolami et al. 2003). Therefore, the findings might not be generalizable to all falls, rather, might be comparable to - some degrees - only with falls with similar perturbation mechanisms. For this very reason, there is a need to have various perturbation and falling replication paradigms for laboratory investigations.

There are three broad categories of inducing an experimental fall: (a) applying an external force/torque to the torso, (b) using moving surfaces to perturb standing balance and (c) self-generating perturbation such as moving body parts. None of these methods simulate an actual fall and hence, do not reproduce the exact fall recovery mechanisms. The problem of “awareness” about the imminent loss of balance is highly pronounced in self-generating perturbations and triggers feedforward and open loop anticipatory mechanisms for preventing the fall (Bouisset and Zattara 1988, Wider et al. 2020). This method is, therefore, mainly applicable for understanding preventative and anticipatory mechanisms in response to loss of balance but might not be sensitive for spontaneous over-ground falling. Moving surface protocols closely replicate falling on slippery surfaces but the results might not be generalizable to spontaneous loss of balance due to aging (Vrieling et al. 2008). It is worth noting that moving the surface perturbs the whole body and causes inertia changes in the Center of Mass (COM). Therefore, there are a cascade of response mechanisms to this type of perturbation (Tang et al. 1998). In biomechanical terms, this type of perturbation induces both translational and angular perturbation to the COM. This paradigm, therefore, activates a combination of balance-restoring strategies.

Impacting the torso causes a sudden displacement of the COM over the base of support (BOS). However, not all falls are initiated by the displacement of COM. This paradigm also causes the COM and center of pressure (COP) to move in the same direction. LOB can be accentuated by the opposite direction of movement of COP and COM. Likewise, in real life situations, not all falls occur by impacting the torso with a mass in a pendular manner. A common feature among all these methods is that all the protocols intend to bring – unexpectedly – the COP to the limits of BOS. The difference is mainly in the level of response activation.

Therefore, different methods provide insights on distinct aspects of loss of balance and falling. A clear understanding of the mechanism of falls would inadvertently provide us with ques for new treatment methods and possibly predicting risk of falling.

Here we introduced and tested a new approach and method for examining loss of balance. The core idea is to elicit reflexive responses (H-reflex) in upright standing, causing contraction in postural muscles and examining the behavior of COP movement along with neuromuscular responses. The loss of balance and falls is inherently a multi-faceted event; and this method can potentially provide new information and biomarkers for certain falling mechanisms. Taken together with other approaches in the literature, this method may prove useful in understanding the complexities of fall risk.

## Method

### New approach

The method entails eliciting the H-reflex in upright standing, causing a single twitch in soleus-gastrocnemius muscles. Soleus H-reflex protocol has been explained in detail in (Knikou 2008) and its functional relevance has been well-documented (Misiaszek 2003, Nematollahi et al. 2017, Tahayori and Koceja 2019, Tahayori, Port, et al. 2012). Briefly, a short duration pulse is applied on the posterior tibial nerve, stimulating sensory and motor fibers resulting in a direct motor response (upon stimulation of motor fibers) and a spinal response (upon stimulation of sensory fibers, mainly Ia fibers). By increasing the stimulation intensity, the H-reflex amplitude decreases due to collision of the impulses inside the motor fiber and eventually a maximum muscle response (M-Max) appears with no H-reflex (Tahayori et al. 2010).

An M-max in Gastrosoleus muscle in standing causes a plantarflexion in the ankle joint and hence a perturbation in standing posture. This type of perturbation is caused by a sudden postural muscle twitch and therefore, can be regarded as an internal perturbation. Unlike applying a mass or a push near to the center of mass of subjects, this perturbation is close to the hinge of the inverted pendulum model, simulating an ankle strategy for balance corrections. COP and force data are good indicators for quantifying the perturbation.

The stimulation, and therefore, perturbation, can be applied unilaterally on one leg or bilaterally on both legs at the same time.

This method, therefore, needs a nerve stimulation device, a forceplate, a surface EMG system and a data collection hardware and software.

### Analysis approach

By applying this method, a clear deflection in COP movement appears after each stimulation. The initial deflection peak is usually followed by a series of smaller amplitude deflections in COP movement. These oscillatory movements tend to bring the system back to its normal equilibrium (Krebs et al. 2001). This pattern of a large initial deflection followed by a train of small corrections is a manifestation of a closed-loop control of posture (Deliagina et al. 2006). As such, the analysis of a normal response provides features for comparison with pathological conditions. A sampler COP deflection pattern in response to stimulation is shown in figure 1.

**Figure 1.**
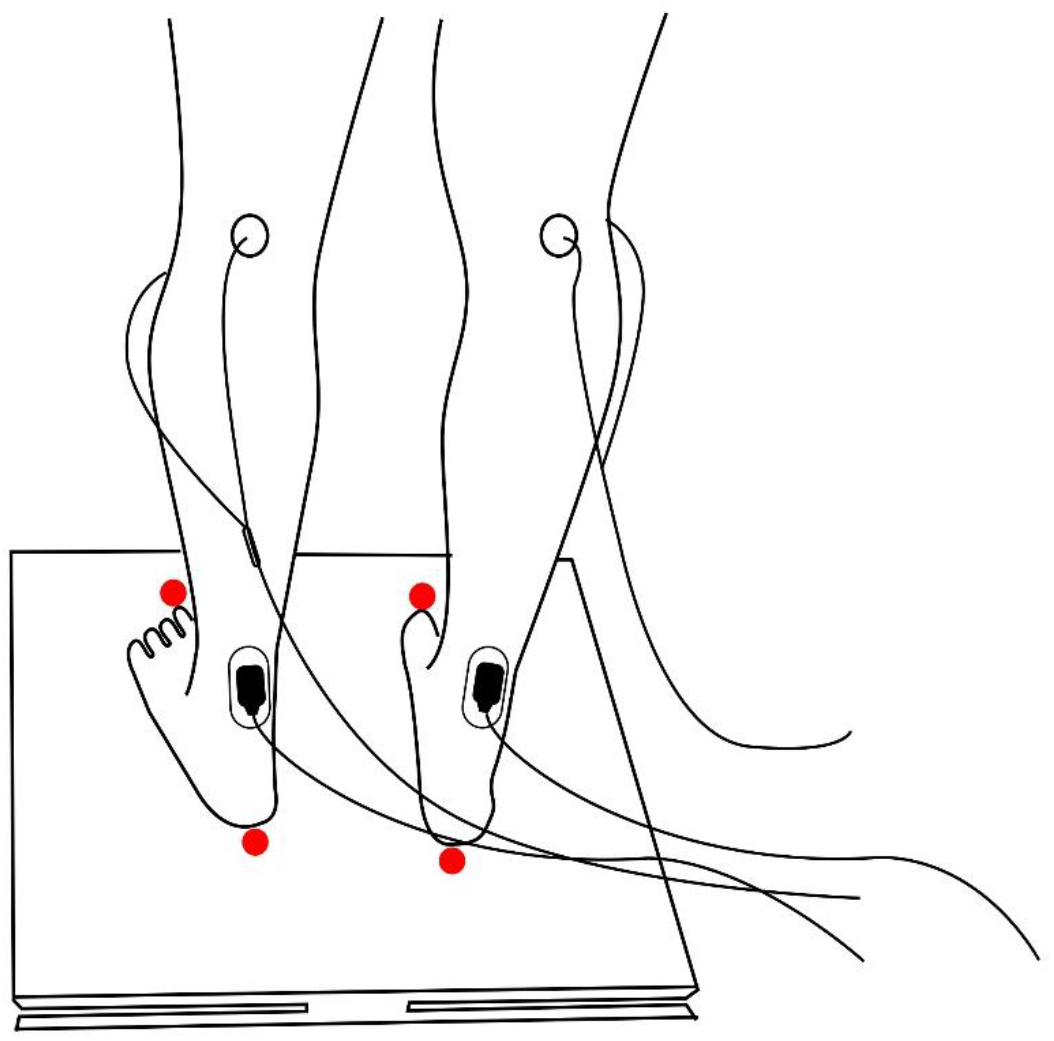
Protocol setup. Stimulation electrodes were placed on the posterior tibial nerve to elicit an M-wave and or H-reflex. The response was recorded by surface electrodes placed on the soleus muscle. The resultant muscle contraction would induce a plantar flexion moment in a closed kinematic chain, causing a perturbation in standing posture. Unilateral or bilateral stimulation would provide sideway or forward perturbations. Red dots represent the places marked for measuring BOS and feet locations.

To quantify the duration of perturbation and determine the instant at which COP movement became stable again, we used a method inspired by the Rambling-Trembling hypothesis (Tahayori, Riley, et al. 2012, Zatsiorsky and Duarte 2000). Our assumption is that these deflections are occurring to bring the COP back to its normal equilibrium points and for that reason, several corrections from the CNS are being applied by moving the COP back and forth (small amplitude overcorrections following a large size perturbation). Therefore, there expects to be a change in the number of equilibrium points in the perturbation zone. The equilibrium points were determined from the following formula:

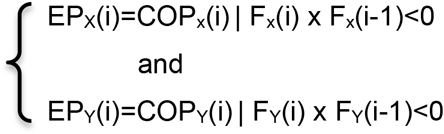

Where EP is the Equilibrium Points, COP is the center of pressure and F is horizontal force from the forceplate. EPs for Anterior-Posterior (AP) and mediolateral (ML) directions were calculated separately.

The density of EPs was measured in the unperturbed trial and an average of EP per second was calculated. EP density with bins of 1 sec duration with a 500 ms overlap window was calculated after each perturbation. The termination of the perturbation was determined by the point at which EPs got back to 85% of the value calculated in the unperturbed period.

Each perturbation can be quantified as a vector with the length of the vector being the size of COP displacement and the angle of the vector being the angle of deviation of the COP. To calculate these vectors, the initial point of the vector was regarded as the average location of COP for 2 seconds prior to the instant of stimulation delivery and the endpoint was regarded as the peak deflection (turning point) of the COP.

The angle of vectors is affected by the orientation and foot placement of the subject on the forceplate. To correct for this confounding factor, feet locations were marked on the forceplate and after the subject stepped off the plate, the placement of these marks were pressed by a metal rod, one at a time, and recorded on a separate file. These points were used to define feet angle. Angle bisectors of the feet angles were calculated and the perturbation vectors were rotated relative to the angle bisector.

The vectors can be resolved to two orthogonal components, corresponding to ML and AP trajectory of COP during perturbation. We have observed that this correction makes minute changes in the results but would be significant if the subjects stood with some angles relative to the coordinate systems of the forceplate.

### Testing protocol

To test this approach and the suggested analytical methods for posture perturbation study, we conducted a study to show how this method yields useful information regarding the mechanism of falling. Fifteen healthy young individuals (age: 26.1±3.2, 9 males, 6 females) who reported to be free of any orthopedic, visual, or neurological disorders participated in this study. This study was approved by the Institutional Review Board (IRB) of Indiana University Bloomington. All subjects signed the written informed consent form prior to participating in the study. Subjects stood barefoot on an AccuSway forceplate (AMTI Inc, Watertown, MA, USA) assuming a normal posture and looked at a target 3 meters in front of them, at their eye level. An initial 60 second unperturbed trial was recorded to measure normal movement of COP with no perturbation.

The H-reflex was elicited while subjects were standing normally on the forceplate. Fifteen stimulations were delivered with minimum intervals of 16 seconds (20 sec ± 4 sec) to ensure that the subjects had fully recovered from perturbation. Stimulations were given either to the right, left or both legs simultaneously.

The electromyography (EMG) activity of the Soleus and Tibialis Anterior (TA) muscles of both legs were recorded using surface EMG electrodes (Therapeutic Unlimited, Company dissolved). Electrodes were placed on the bulk of the muscles, parallel to muscle fibers.

Stimulations were delivered through a DS7A constant current stimulator (Digitimer, Welwyn Garden City, Hertfordshire, UK) with a pulse width of 1 ms. Active stimulation electrode was placed at the back of the knee on the posterior tibial nerve. Passive electrode was placed on the knee cap. The intensity of the pulses were increased until a saturated M-response (M-max) was observed. Care was given as to avoid contamination of the fibular nerve (Tahayori et al. 2015). This was verified by a strong contraction of Gastrosoleus and no contraction in the peroneal muscles.

Forceplate, EMG data and stimulation timestamps were fed into an Analog to Digital Converter (NI-6211-USB, National Instrument, Austin, Texas, USA) and digitized at 4000 samples per second. A customized program written in DASYLab (Measurement Computing Corporation, Norton, MA, USA) was used to observe the signals, measure M-max values in real-time and save data on a computer for further offline analysis.

Figure 1 is an illustration of the setup, showing the standing position with stimulation and recording electrodes in place.

The COP signal was calculated using the conventional method provided by the manufacturer. The instants of perturbation were extracted from the timestamps of the stimulator machine. A part of the offline analysis were performed in custom-written programs with MATLAB R2020a (MathWorks, Natick, MA, USA). Certain analyses were performed in Python.

## Results

The electrical stimulation of the posterior tibial nerve elicited an M-max in the corresponding Soleus muscle and induced a perturbation in subjects’ posture. This was manifested by a sudden deflection of COP trajectory. A sampler EMG response including M-max and subsequent EMG activity is shown in figure 2.A. To better observe the EMG signal after the artifact caused by stimulation, the signal was scaled to uV and is shown in the inset of panel A. COP deflection as a result of this M-wave is presented in figure 2.B.

**Figure 2.**
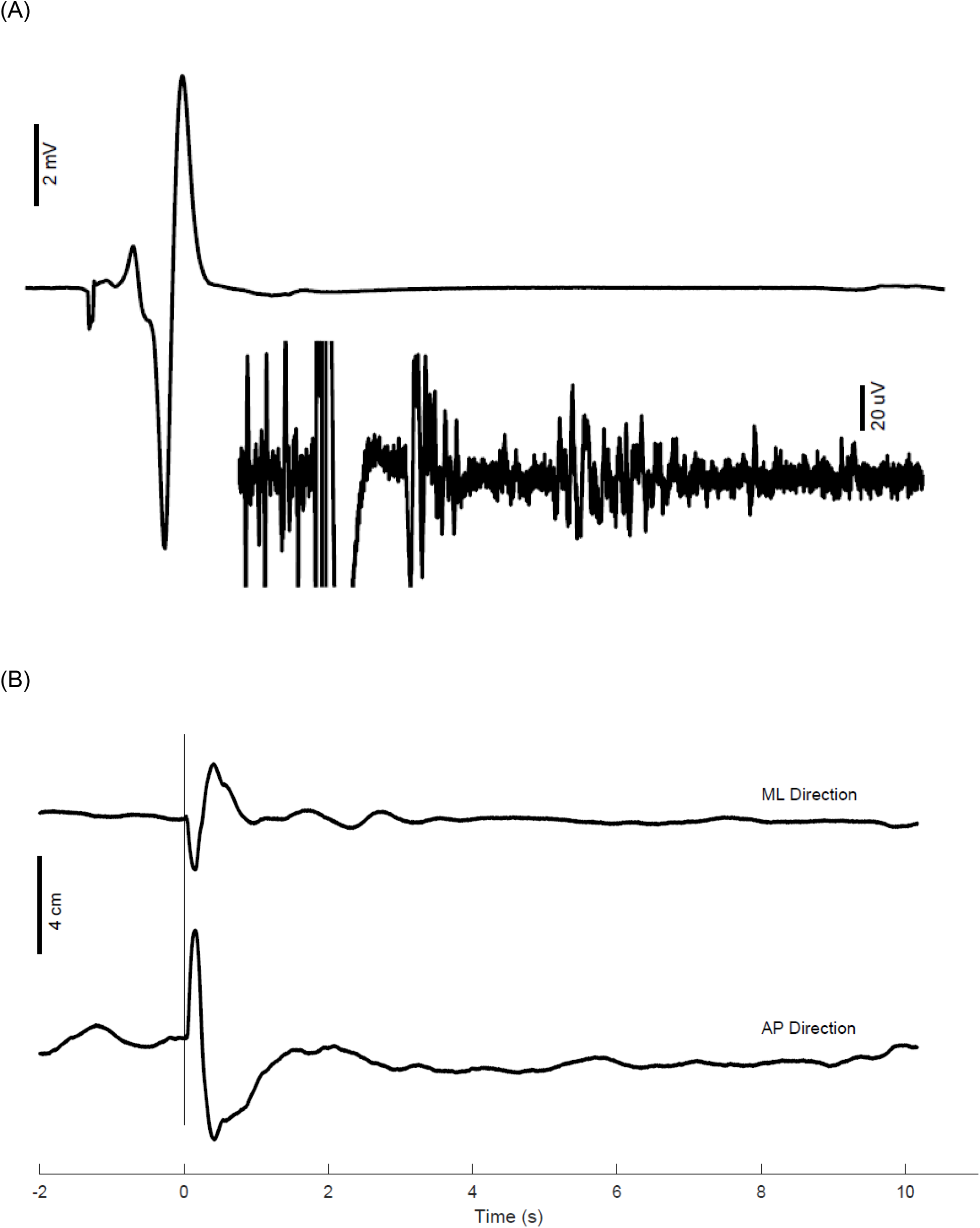
(A) Sample M-max with almost no observable H-reflex. The inset is an amplification of the same signal to show EMG activity, (B) Sample of COP trajectory deflection.

The amplitude of M-max was monitored after each stimulation to prevent any significant change in the response and inconsistency in the perturbation caused by this stimulation. Figure 3 shows the peak-to-peak amplitude of M-waves in a sampler subject, showing a normal fluctuation in the amplitude but no substantial drift.

**Fig 3.**
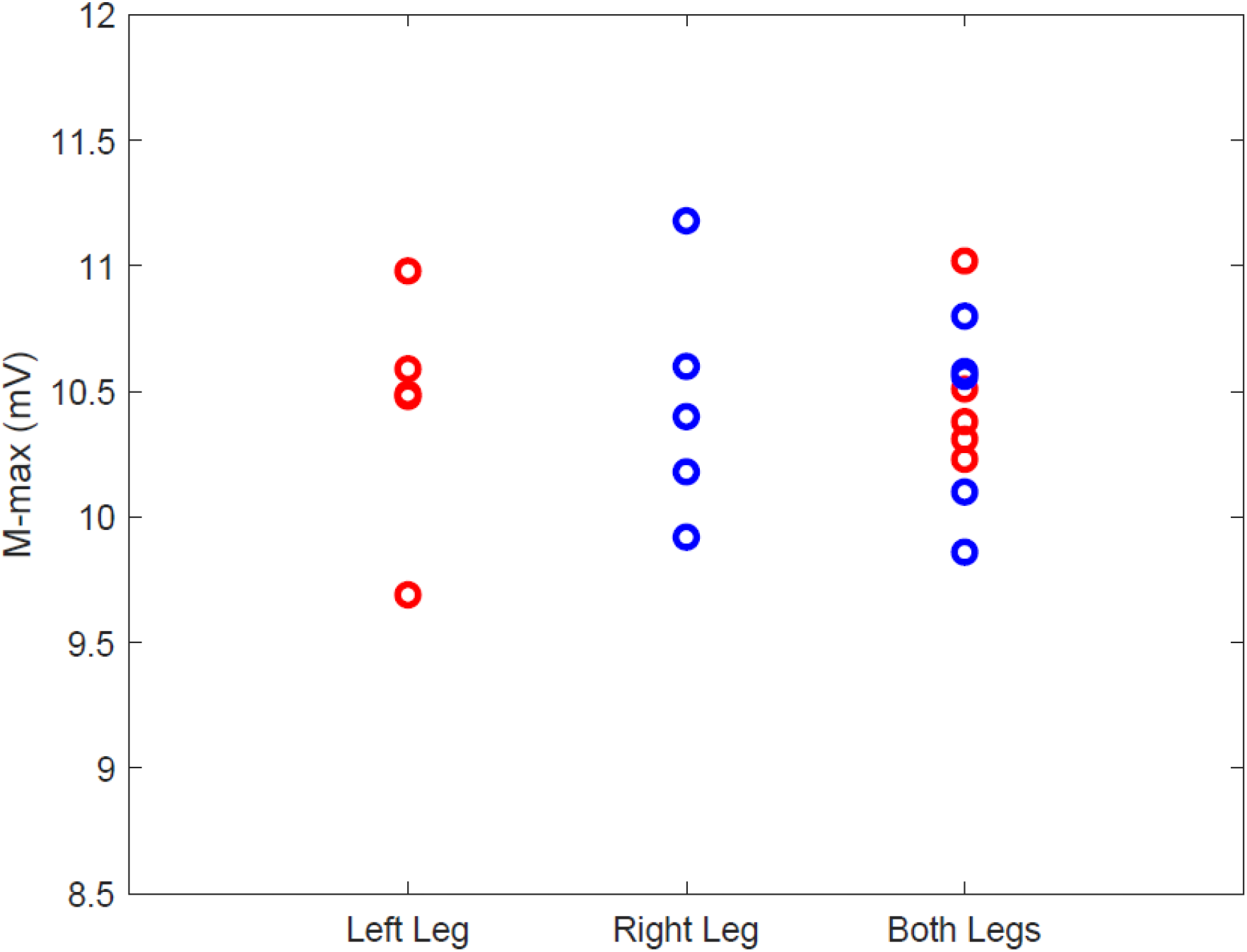
M-max consistency. The amplitude of M-max was monitored for both lower extremities.

### Equilibrium points

The density of Equilibrium Points (EPs) were calculated in the perturbation period. There was a significant difference in the side of stimulation with least density of EP in bilateral stimulation and largest number in right leg stimulation. The results are shown in figure 4.A. The duration of stimulation was calculated based on the return of EP density to 85% of normal values. The duration was significantly longer for bilateral stimulation condition. The results are shown in figure 4.B.

**Fig 4.**
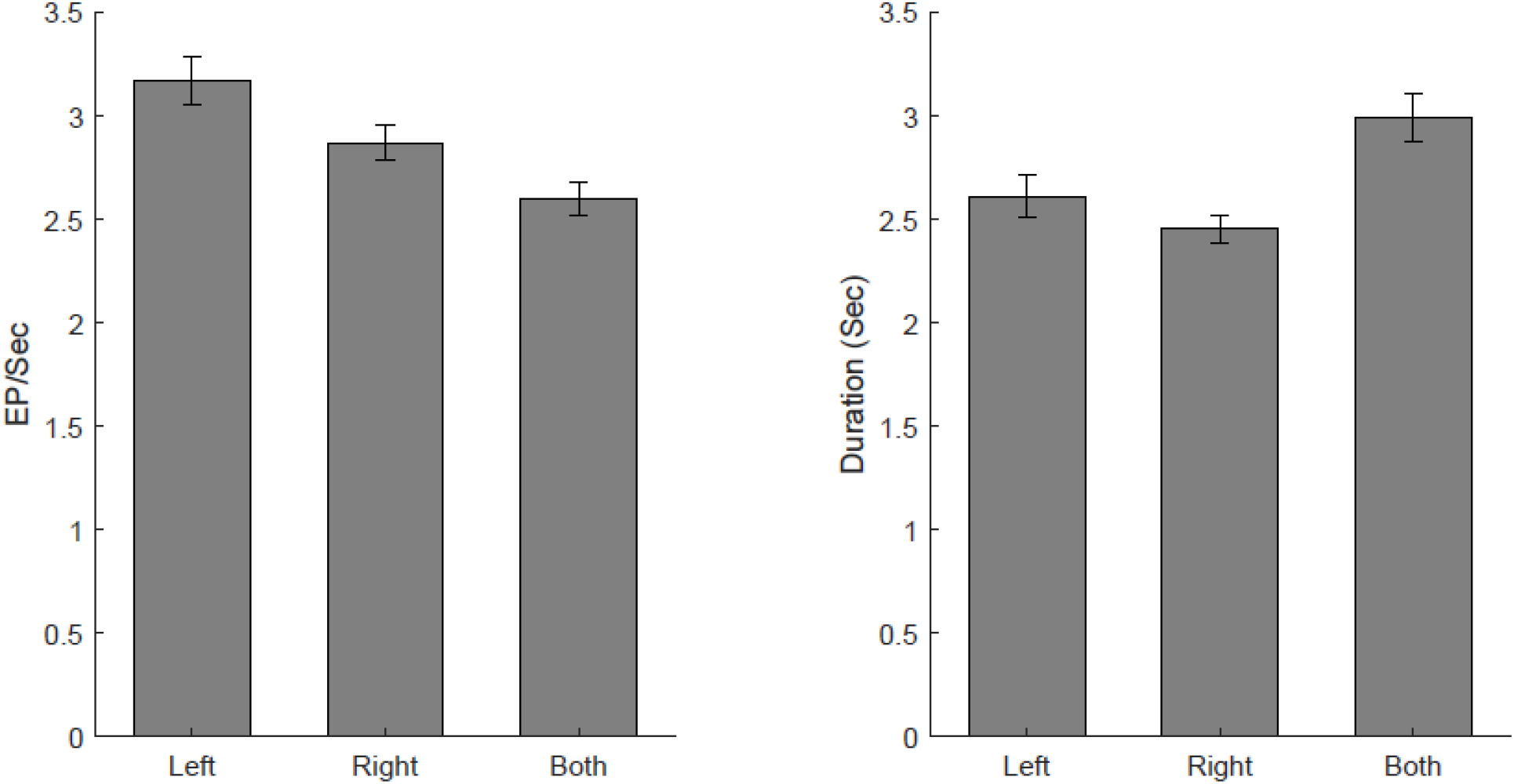
(A) Density of Equilibrium points, (B) Duration of perturbation.

### Vector analysis

The initial deflection in the postural sway was presented as a forward-direction vector. The size of the vector is an indicator of perturbation amplitude. The direction of the vector is an indicator of how the COP moved in response to perturbation. Figure 5.A shows a sample of these vectors from a top view (looking from above over the forceplate area)

**Fig 5.**
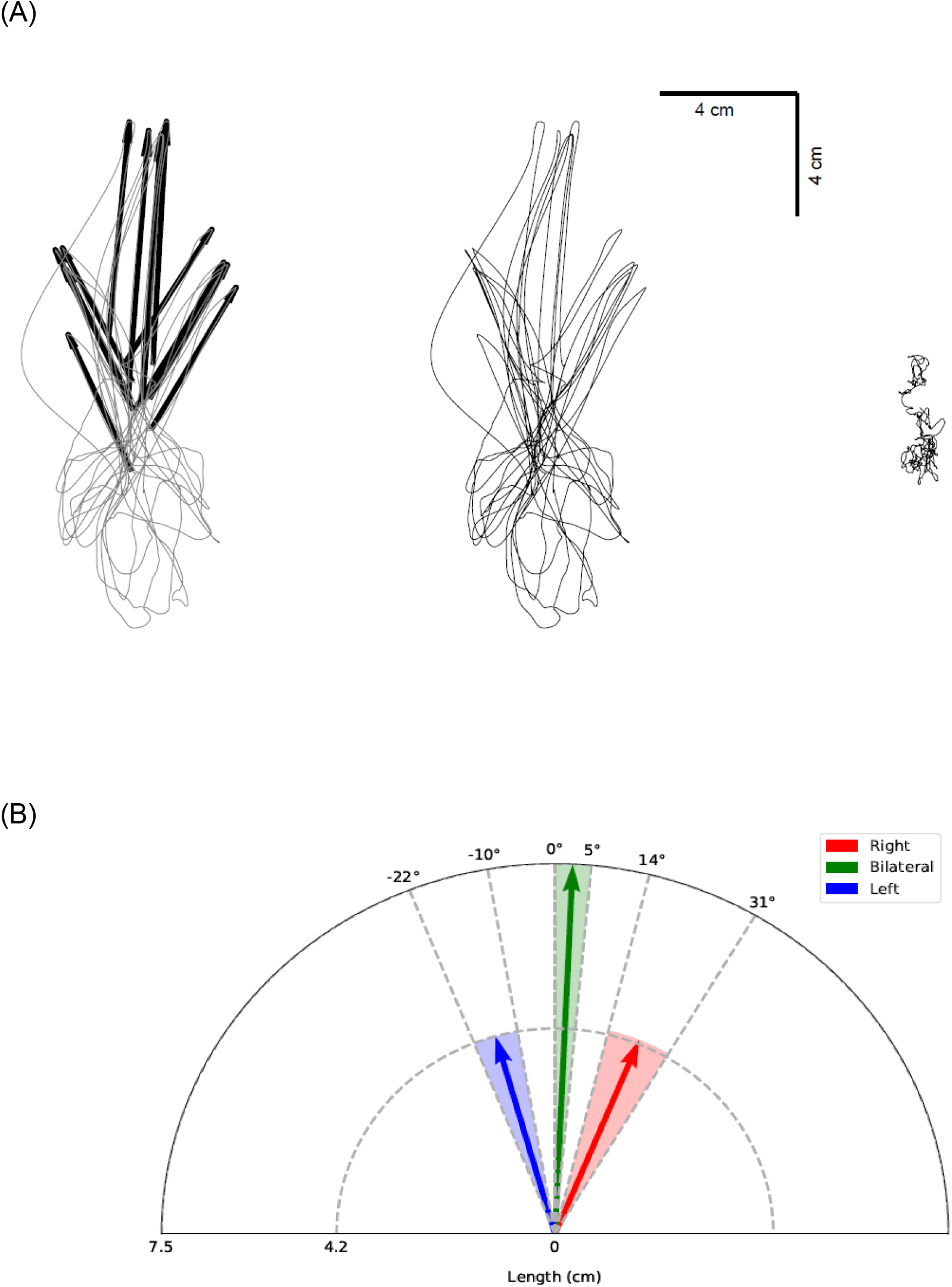
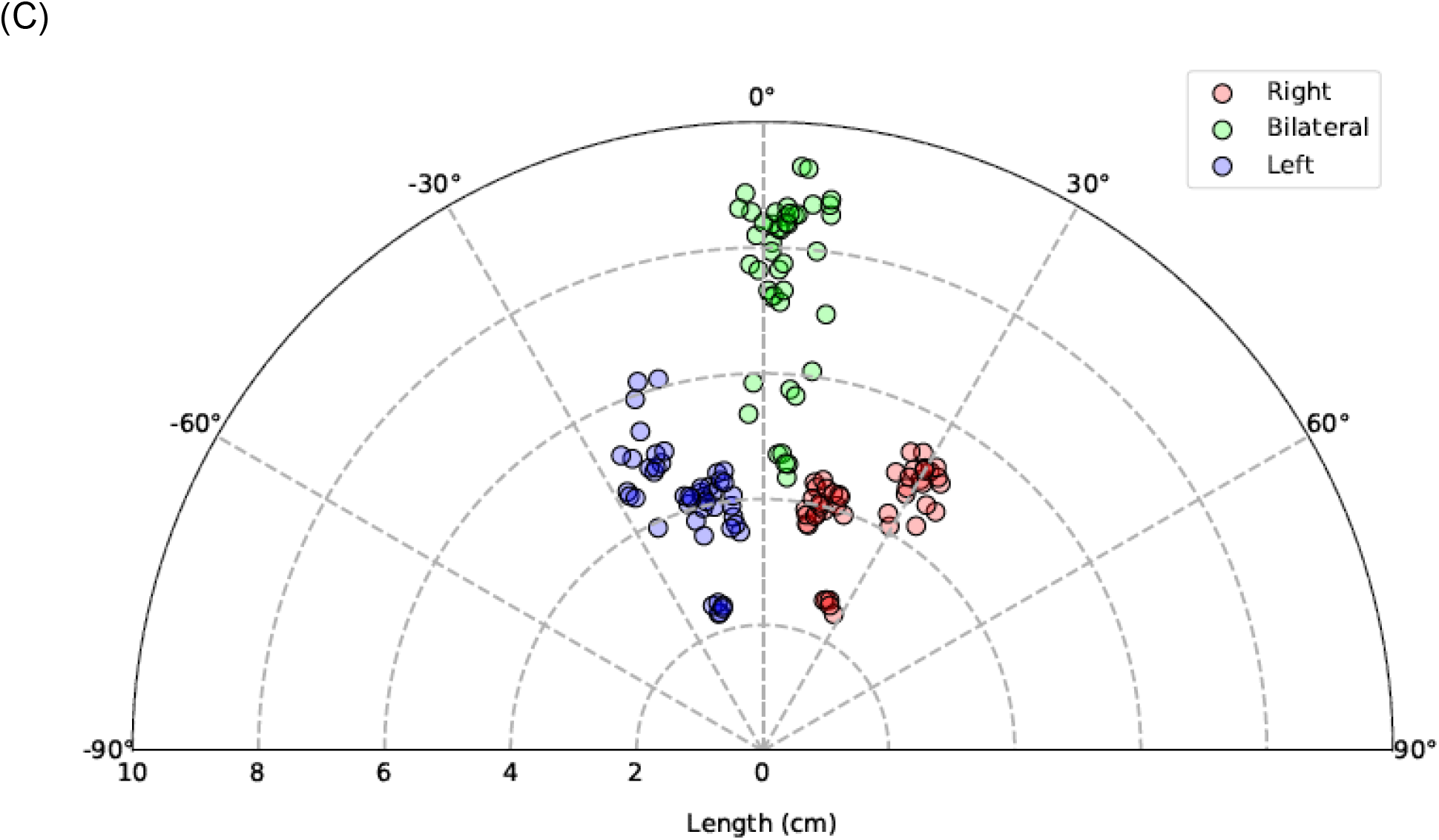
Vector analysis of perturbation. (A) Sample of a typical subject. From left to right: COP perturbation with vectors overlayed. COP perturbation without vectors (For clarity), unperturbed COP. (B) Vector analysis results for all subjects is shown in this graph. The shaded area represents the SD of each perturbation direction with vectors representing the direction and amplitude of perturbation. (C) Same as in figure B but instead of vectors, actual points are plotted. A line connecting the center (0,0) to each point would constitute its vector.

The three distinct directions of vectors, corresponding to the side of stimulation, are clearly observable. The results are summarized in figures 5.B and 5.C.

## Discussion

Simulation of loss of balance and falling is a challenging laboratory experiment. Normal and skillful movements are relatively easy to replicate in labs, given that there is always a laboratory effect to change motor behavior (Tamburini et al. 2018). However, examining an unskillful movement with an unpredictable nature is moving to a new level of difficulty in lab-based movement analysis. Therefore, in laboratory settings, a paradigm is defined to artificially trigger LOB response mechanisms and measure the reactions, or lack thereof.

Sliding surfaces displace the whole body in a closed kinematic chain, thereby leaving the COM behind and causing loss of balance. Impacting the torso displaces the COM and might or might not be accompanied by hip and stepping strategies. Angular Momentum Perturbator (AMP) paradigm is a sophisticated method to induce angular torque without terribly affecting COM movement (Schumacher et al. 2019). Self-generated perturbation strategies intend to use body parts movement to induce perturbation in upright standing. Sudden upward movements of arms is such an example. However, this type of perturbation is accompanied by feedforward reactions and entails anticipatory postural adjustments (Hatzitaki et al. 2005, Kubicki et al. 2012). Therefore, such protocols are useful particularly for understanding the mechanisms of predictable loss of balance. Comparatively mechanical stimulations take longer to apply but the celerity of an electrical stimulation and the resultant muscle twitch makes it more similar to an actual loss of balance.

Galvanic stimulation (GVS) has been used for stimulating the vestibular labyrinth. GVS has been routinely used to study the vestibular system and to simulate temporary vestibular impairment and its effect on locomotion and posture (Bacsi and Colebatch 2005, Deshpande and Patla 2005). does not induce a sudden mechanical perturbation, rather it causes a temporary derangement in the vestibular system, affecting posture. This method has shed light on the influence of the vestibular system on postural control. Depending on the head’s orientation, GVS induces directional perturbation on posture, affecting ML and AP planes differently (Coats and Stoltz 1969, Day et al. 2002, Tax et al. 2013). This is a similarity of GVS with our method. However, although GVS perturbs COP trajectory, to the best of our knowledge it is not being used to cause a sudden perturbation in balance and simulating a fall.

The method presented here uses a purely artificial reflex protocol to cause muscle twitch and therefore, perturb balance. Ordinarily falls are not caused by a sudden, unexpected twitch of postural muscles. The brief twitch in postural muscles at the ankle joint (the hinge joint in the inverted pendulum model), is the origin of perturbation as well as the first line of response to perturbation (Winter et al. 1990). As such, monitoring and controlling the amount of muscle contraction is easily attainable in this method. Being a closed loop perturbation-adjustment mechanism, this method is suitable for neuromechanical studies of posture and perturbation. This method, in all likelihood, triggers a less complicated control mechanism (rather than triggering multiple levels of responses) which is another strength of the method. More importantly, electrical Stimulation of posterior tibial nerve causes a contraction in the biarticular Gastrocnemius muscle. The role of biarticular muscles in control of posture and response to perturbations have been emphasized over the past few years. However, these muscles are not substantially active in the sliding surface and push-pull paradigms (Schumacher, Berry, Lemus, Rode, Seyfarth and Vallery 2019).

Inducing localized or generalized fatigue is another method of examining postural perturbation. There are examples in the literature to induce fatigue in postural muscles and observe the changes in postural sway (Corbeil et al. 2003, Davidson et al. 2009, Kennedy et al. 2012). This approach follows the same principle of inducing a change in the sensory-motor system and measuring subsequent alterations in its dynamics. The introduced approach here can be incorporated with muscle fatigue to investigate not only the effect of muscle fatigue but also that of the neural fatigue in alterations in postural responses.

We have provided a new method for inducing an internally generated perturbation. Here we used M-max for causing a maximum muscle twitch and largest possible perturbation. It is quite possible to use H-max for inducing perturbation. Muscle contraction caused by H-max is not as strong as that of the M-max but has the advantage of being a neural signal which is known to be modulated as a part of motor programming and sensory-motor interaction (Tahayori and Koceja 2012). Therefore, our method not only provides biomechanical markers for quantifying perturbation, but also neural insights about motor programming at the level of final common pathway. This method does not claim any superiority over the already established methods or claims to solve inherent difficulties associated with the study of falling/loss of balance in a laboratory setting. Rather, it provides meaningful neuromechanical biomarkers with the potential for predicting risk of falling in different populations including neurological patients and elderly people. This method also has potential uses in clinical trial intervention studies. By controlling the degree of internal perturbation to the neural system, changes in recovery can be accurately determined.

As a proof of concept, we conducted a study using the method on a group of healthy young individuals. As was expected, the amplitude of muscle contraction was consistent among trials (as shown by a stable M-max -figure 3) and the three different directional perturbations (to the left, right and both) were clearly observable with distinct and meaningful angle of perturbation (figure 5).

Falls caused by mechanical disturbances can be divided into two wide categories: Base of Support (BOS) falls (Topper et al. 1993) and Center of Mass (COM) falls (Maki and McIlroy 1996). BOS falls are mainly associated with sensory input deficiencies, lower leg muscle weakness and lack of coordination. COM falls mainly occur due to sudden impacts on the trunk, muscle weakness of trunk and sudden body parts movements which make the COM displacement without timely postural adjustments (Voermans, Snijders, Schoon and Bloem 2007). Falls due to peripheral neuropathy, generalized weakness, and drop foot are most likely follow the BOS mechanism while Parkinson’s loss of balance is mainly due to BOS mechanisms. Our method is most probably more suitable for investigating BOS mechanisms while Self-induce perturbations and trunk impact methods are more suitable for investigating COM mechanisms.

In summary, this paradigm provides a new method for investigating mechanism of falls and has the ability to induce unilateral or bilateral perturbation. The method does not use any external perturbation source and has a faster time scale than mechanical perturbations. Future studies are needed to define the limits of normal values based on age.

## Acknowledgement

We would like to thank Ms. Shabnam Hosseini for drawing the illustration. The first author was funded by a predoctoral fellowship from American Heart Association (13PRE14780094) and sincerely thanks the Association for this award and funding.

